# Membrane association is required for Stmn2-mediated axon protection

**DOI:** 10.1101/2022.12.23.521801

**Authors:** Emma J.C. Thornburg-Suresh, Jerianne E. Richardson, Daniel W. Summers

## Abstract

Axon integrity is essential for functional connectivity in the nervous system. The degeneration of stressed or damaged axons is a common and sometimes initiating event in neurodegenerative disorders. Cellular factors that preserve axon integrity have an important influence on the fate of a damaged axon. Stathmin-2 (Stmn2) is an axon maintenance factor that is depleted in Amyotrophic Lateral Sclerosis and replenishment of Stmn2 can restore neurite outgrowth in diseased neurons. Stathmins have a well-documented role in microtubule dynamics during neurodevelopment, yet mechanisms responsible for Stmn2-mediated axon maintenance in injured neurons are not known. We demonstrate that membrane association of Stmn2 is critical for its axon-protective activity. Axonal enrichment of Stmn2 is driven by palmitoylation as well as tubulin interaction. We discover that another Stathmin, Stmn3, co-migrates with Stmn2-containing vesicles and undergoes regulated degradation through DLK-JNK signaling. The Stmn2 membrane targeting domain is both necessary and sufficient for localization to a specific vesicle population and confers sensitivity to DLK-dependent degradation. Our findings reveal a broader role for DLK in tuning the local abundance of palmitoylated Stathmins in axon segments. Moreover, palmitoylation is a critical component of Stathmin-mediated axon protection and defining the Stmn2-containing vesicle population will provide important clues toward mechanisms of axon maintenance.

## INTRODUCTION

Neurons extend long structures called axons that establish connections throughout an organism. In humans, some axons project over a meter in length to reach their targets. This incredible distance poses unique biological challenges on the axon compartment. One such challenge is the transport of cargo through the entire length of the axon to supply nerve terminals with proteins necessary for function. Axonal damage or stress can impair transport and threaten axon integrity. For example, axon transection deprives distal axon segments of the survival factor Nmnat2 and stimulates Sarm1-mediated axon dismantling (1,2). Loss of axon health is an early sometimes initiating event in a variety of neurodegenerative disorders in the peripheral and central nervous systems (3). Therefore, defining cellular pathways that control axon susceptibility to pathological degeneration is an important need.

The microtubule-binding phosphoprotein Stathmin-2 (Stmn2) is an axon maintenance factor that is depleted in Amyotrophic Lateral Sclerosis (ALS). ALS-linked mutations in TDP-43 cause aberrant splicing of Stmn2 mRNA and decreased Stmn2 protein expression in humans with familial ALS (4-6). In mouse models, loss of Stmn2 provokes motor and sensory neuropathies (7-9), and selective knockdown of Stmn2 in dopaminergic neurons leads to neuron loss and locomotor deficits (10). Moreover, replenishing Stmn2 restores neurite outgrowth in iPS-derived motor neurons from patients with ALS (5). These discoveries collectively reinforce the important contribution of Stmn2 to axon maintenance.

Stmn2 is a member of the Stathmin family of phosphoproteins that include Stmn1, Stmn3, and Stmn4 (11). Proteins in the Stathmin family directly bind tubulin heterodimers to regulate microtubule stability. Stathmin interaction with tubulin heterodimers is regulated by phosphorylation events in an unstructured proline-rich domain (PrD) present in all Stathmin proteins (12). For example, the MAP Kinase JNK phosphorylates Stmn2 on serines in the PrD and thereby regulates microtubule dynamics in neurons (13,14). However, phosphorylation can also impact Stmn2 function beyond tubulin interaction. JNK-mediated phosphorylation of Stmn2 promotes regulated degradation of this protein (15). Alanine substitutions in two serines phosphorylated by JNK boosts local abundance of Stmn2 in axons and overexpression of this phospho-dead Stmn2 variant delays axon degeneration in transected axons (15). JNK is phosphorylated downstream of the MAP3K Dual Leucine Zipper Kinase (DLK) and stimulating DLK accelerates loss of Stmn2 protein from axons (16). Whether the DLK-JNK signaling pathway similarly regulates degradation of other Stathmin family members is unknown.

In addition to phosphorylation, Stmn2 is palmitoylated on two cysteine residues in a N-terminal membrane targeting domain (MTD) (17). The Stmn2 MTD is necessary and sufficient for localization to the Golgi as well as vesicles in the growth cone (18,19). Recombinant fragments of Stmn2 lacking the MTD can still bind tubulin heterodimers and regulate microtubule stability *in vitro* (20). Therefore, membrane association is not necessary for microtubule binding. Palmitoylation is required for Stmn2-dependent regulation of amyloid precursor protein processing and trafficking to the cell surface (21), indicating an important contribution to Stmn2 function in neurons. However, a role for Stmn2 palmitoylation in axon degeneration has not been examined.

We discovered that palmitoylation is required for Stmn2-mediated axon protection. Blocking Stmn2 palmitoylation did not prevent Stmn2 from accumulating in axons as the tubulin binding region also contributes to axonal localization. A highly related MTD is present in Stmn3 and functionally compensates for the corresponding MTD in Stmn2. Moreover, Stmn2 and Stmn3 co-migrate on the same vesicle population. Consistent with behavior observed for Stmn2, Stmn3 is short-lived, rapidly lost from transected axons, and undergoes regulated degradation in response to DLK-JNK signaling. Notably, Stmn1 lacks a membrane targeting domain and does not display these features. The Stmn2 MTD is sufficient to localize a reporter protein to Stmn2-containing vesicles and confer DLK-dependent instability in axon segments. Collectively, these observations identify new functional roles for membrane association in Stmn2-mediated axon integrity.

## EXPERIMENTAL PROCEDURES

### Reagents and Plasmids

Human Stathmin expression clones were from Genscript (STMN1-OHu14092D, STMN2-OHu14465D, STMN3-OHu10050D) and Gibson cloning was used to generate lentiviral expression plasmids. All lentiviral expression plasmids are under control of the human ubiquitin promoter. Phenol-red free neurobasal, B27 supplement, L-glutamine, penicillin-streptomycin, and Opti-mem were from Gibco. HEK293T cells were cultured in DMEM (4.5g/L glucose; Corning) supplemented with heat-inactivated Fetal Bovine Serum (Corning), 2mM L-glutamine, and penicillin/streptomycin (10U/mL). Western immunoblotting was performed with the following primary antibodies; anti-GFP (Thermo Fisher; RRID:AB_221569; 1:1,000), anti-Stmn1 (Cell Signaling; RRID:AB_2798284; 1:1,000), anti-Flag (Cell Signaling, RRID:AB_2572291; 1:1,000). anti-Stmn2 (R&D Systems; RRID:AB_10972937; 1:1,000), anti-Stmn3 (Proteintech; RRID:AB_2197399; 1:1,000), anti-Tuj1 (Biolegend; RRID:AB_2562570; 1:10,000). Detection of primary antibodies was performed with the following secondary antibodies that were visualized with a LI-COR Odyssey imager (Thermo Fisher; RRID:AB_1965956 and RRID:AB_2556622). GNE-3511 and JNK inhibitor VIII were from Cayman Chemicals.

### Isolation and Culture of DRG Sensory Neurons

Embryonic sensory neurons derived from dorsal root ganglia (DRG) were isolated from embryonic day (E) 13.5 mouse pups (equal number of male and female embryos). For all studies, pregnant CD1 mice were purchased from Charles River Laboratories. DRG sensory neurons were dissociated with trypsin and seeded as droplets in tissue-culture treated plastic dishes pre-coated with poly-d-lysine (PDL) and laminin. DRG neurons were cultured in phenol red free Neurobasal Medium supplemented with 2mM glutamine, 10 U/mL penicillin/streptomycin, 2% B27 supplement, 50ng/mL nerve growth factor, and 1μM 5-fluoro-2’-deoxyuridine/ 1μM uridine to inhibit growth of non-neuronal cells. Fresh culture media was added every 2-3 days. For axon degeneration experiments, DRG neurons were infected 2 days after plating (DIV2) with lentiviral particles expressing myristoylated-mRuby3 (myr-mRuby3) to label axon membranes. DRGs were also transduced with lentiviruses that express Bcl-xL to prevent nonspecific toxicity from the overexpression of our lentiviral constructs. Bcl-xL overexpression does not affect the kinetics of axon degeneration in Wallerian degeneration (22). On DIV2-4, lentiviruses containing Stathmin expression constructs were applied to neurons. All experimental protocols in this study using mice were reviewed and approved by the University of Iowa Office of the Institutional Animal Care and Use Committee.

### Lentiviral Preparation and Transduction

To produce lentiviral particles, HEK293T cells were transfected with plasmids expressing vesicular stomatitis viral G protein, the lentiviral packaging plasmid psPAX2, and a lentiviral expression plasmid derived from the FCIV backbone (ubiquitin promoter) (23). On DIV2-3, media containing the lentiviral particles was collected, centrifuged for 1min at 500 x g to remove debris, and the supernatant used to transduce DRG cultures on DIV2-4.

### Measurement of Axon Degeneration

Axon degeneration was measured as previously described (24). For axotomy studies, DRGs were seeded in 24 or 96-well plates. On DIV6 or DIV7, axons were manually severed from the cell body with a razor blade. Fluorescent images of severed axons were acquired with the Cytation 5 automated microscope (Agilent). Within one experimental replicate, at least three wells were assessed per condition, within each well six independent fields were collected. Independent DRG sensory neuron cultures are used for each experimental replicate. For time course studies, the same axon fields were imaged for the duration of the timeseries. Axon degeneration was scored from a custom-generated ImageJ macro (24) that scores fragmented/circular axon segments as a ratio of total axon area for each image. The macro assigns each image a number between 0 – 1 with a higher value representing greater axon degeneration.

### Biochemical Analysis of Axon-Only Protein extracts

For analysis of protein levels in axon-only fractions, DRGs were densely seeded in a spot culture in a 12-well plate coated with PDL/laminin with fresh media added every 2 days. On DIV 7 or DIV 8, a razor blade is used to cut around the cell bodies and the spot of cell bodies is removed with a pipette. Axons were washed with PBS then lysed in RIPA buffer (50mm Tris-HCl pH 7.4, 1mM EDTA, 1% Triton X-100, 0.5% sodium deoxycholate, 0.1% sodium dodecyl sulfate, 150mM NaCl, 1mM phenylmethylsulfonyl fluoride, 1X EDTA-free protease inhibitor cocktail [Genesee Scientific], and phosphatase inhibitors 5mM NaF and 1mM NaVO_4)_. Axon-only extracts were precleared by centrifugation (at 4°C, 2,500 xg for 5 minutes) and then the supernatant was mixed with sample buffer (65.2mM Tris-HCl pH 6.8, 2% SDS, 10% glycerol, 8% beta-mercaptoethanol, 0.025% bromophenol blue). The samples were boiled before analysis by SDS-PAGE and western immunoblotting. For analysis of protein turnover, the DRG neurons are treated with 25ug/mL cycloheximide to inhibit protein synthesis. Cycloheximide was added on DIV 6 or DIV 7. The time points for cycloheximide addition are indicated per-experiment. Lysates were analyzed by SDS/PAGE and Western immunoblotting for the indicated protein and normalized to levels of tubulin using Tuj1. All quantification of Western immunoblotting is performed with ImageJ.

### Comigration Imaging and Analysis

For analysis of comigration, neurons are sparsely plated in 3-4 spots on 35mm plates (Fluorodish, World Precision Instruments) coated with PDL/Laminin. Lentiviruses expressing differentially tagged constructs (Venus or mCherry) were added on DIV3 or DIV4. Cultures that displayed axonal blebbing or other signs of degeneration were not used. Images were collected with a LEICA TCS SP8 Confocal Microscope using a 63X oil immersion lens (N.A. 1.20) and resonant scanner. The 35mm plates are placed into an Okolab incubation chamber, and neurons were maintained at 37°C, 5% CO_2_, and 20% O_2_ for the duration of the experiment. Images were collected simultaneously in both the red and green channels as well as brightfield to identify axon segments. At least three independent experimental replicates were completed per condition.

Analysis of comigration is conducted in a manner to ensure the experimenter is blinded to the imaging conditions. Prior to analysis of a mobile puncta, the axon is checked to ensure that both viruses are expressed. A mobile puncta is identified in one of the two fluorescent channels and a kymograph is generated for that particle using the ImageJ Kymograph Builder and merged with a kymograph generated from the corresponding channel. Merged kymographs and individual tracts are scored as comigrating or not. The kymographs are scored for comigration by at least two lab members. Between 75-100 puncta are analyzed per experimental condition. Percentage of comigration with Stmn2 is calculated per-experimental condition.

### Fluorescence Microscopy analysis of Stathmin-Venus levels in axons

DRG sensory neurons seeded in 96-well plates were transduced with lentiviruses on DIV3 to express the indicated Stathmin and myristoylated mRuby3 to label axons. On DIV6, DRGs were fixed in 4% paraformaldehyde and stored in phosphate buffered saline. Fixed axons were visualized with a Leica DMi8 inverted fluorescence microscope using a 20X objective (N.A. 0.4) maintaining equivalent settings during image acquisition across experimental replicates. Images were captured with a Leica DFC7000T and processed with ImageJ. Thresholding of myristoylated Ruby3 axon images was performed to generate axon region-of-interests (ROIs) for each image. We used rolling ball background subtraction on Venus images and we measured fluorescence intensity of the Venus image within the corresponding axon ROI. For each experimental replicate, at least three independent wells were used per condition and at least two fields were collected per well. To measure fluorescence intensity in the soma, we used rolling ball background subtraction on all Venus images then measured mean fluorescence intensity within the soma using a standard ROI.

### Statistics

All statistical analysis was performed with GraphPad Prism. Specific statistical tests applied for each experiment are identified in the corresponding figure legend.

## RESULTS

### Stmn2-mediated axon protection requires palmitoylation

Stmn2 is regulated by two different post-translational modifications, phosphorylation and palmitoylation. Serine-to-alanine replacement of JNK phosphorylation sites (Stmn2AA) enhances Stmn2 axon protective activity (15). Protein levels of Stmn2AA were also elevated due to extended half-life in axon segments suggesting that prolonged stability is responsible for axon protection. Stmn2 protein turnover is also regulated by palmitoylation (25) yet a functional role for this post-translational modification in axon degeneration has not been examined. To address this question, we converted two cysteine residues modified by palmitoylation to serines (Stmn2CS) and evaluated how the loss of palmitoylation impacts Stmn2 activity in Wallerian degeneration, a model of pathological axon loss.

We over-expressed wildtype or mutant forms of Stmn2 tagged with Venus via lentiviral transduction in embryonic-derived, mouse sensory neurons from dorsal root ganglia (DRG). For all experiments, we used a Stmn2 human coding sequence that shares 100% amino acid homology with mouse Stmn2. Three days following lentiviral transduction with Stmn2 expression constructs, axons were cut with a razor blade to induce degeneration of severed axon segments. We replicated past findings that overexpression of Stmn2AA delayed axon degeneration 10-hours post-axotomy (Figure 1A&B). Overexpression of Stmn2CS behaved similar to wildtype Stmn2 and did not affect axon degeneration (Figure 1A&B). To assess whether palmitoylation is necessary for axon protection evoked by Stmn2AA, we generated a version of Stmn2 lacking both JNK phosphorylation and palmitoylation (Stmn2; double dead; DD). In contrast to Stmn2AA, Stmn2DD did not suppress axon degeneration 10 hours post-axotomy (Figure 1A&B), indicating that palmitoylation is required for Stmn2AA-mediated axon protection.

**Figure 1.**
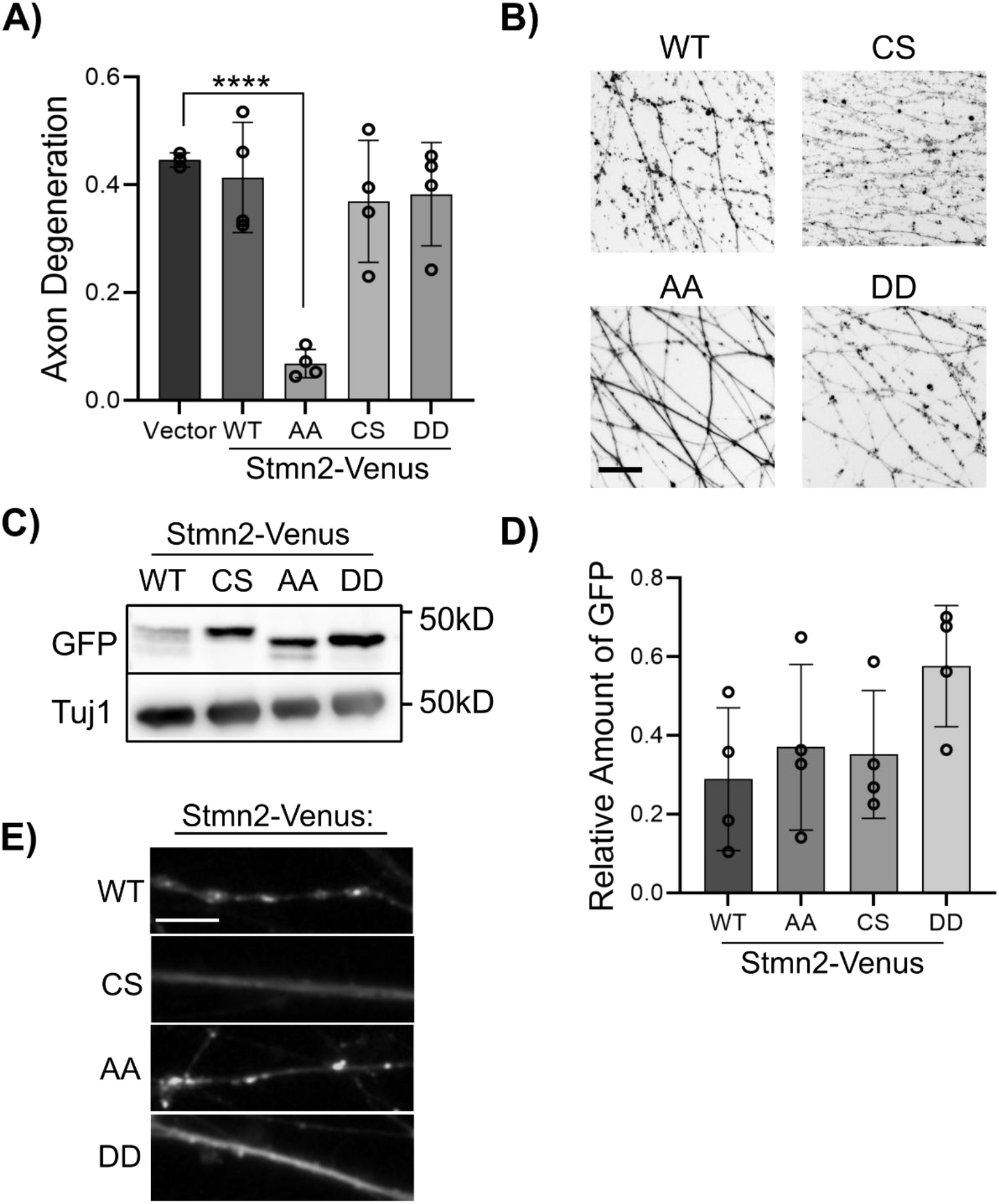
Palmitoylation is required for Stmn2-mediated axon protection. *(A)* DRG sensory neurons were transduced with lentiviruses expressing the indicated human, Venus-tagged Stmn2 constructs and axons were severed with a razor blade. The degeneration of distal axons was measured 10hr post axotomy (WT wildtype; CS C22,24S; AA S62,73A; DD C22,24S and S62,73A). Each Stmn2 construct was compared to the empty vector control (n = 4, vector vs. AA, unpaired t-test, ****p< 0.0001). *(B)* Representative images of distal axons 10hr post axotomy labeled with myristoylated mRuby3. *(C)* Western blot of Venus-tagged Stmn2 variants from axon-only extracts with quantification shown in *(D)* (n = 4). *(E)* Fluorescence microscopy of Stmn2-Venus proteins demonstrating localization to axon segments. Scale bar = 5μm. Error bars represent +/-1 standard deviation (STD).

We suspected that loss of palmitoylation would interfere with axonal transport of Stmn2, which may account for the lack of axon protective activity observed from the Stmn2DD construct. We examined protein levels of each Stmn2 construct in axons by western immunoblotting and fluorescence microscopy. As previously observed (15,25), Stmn2AA and Stmn2CS protein levels were increased in axon-only extracts as compared to wildtype Stmn2. Notably, Stmn2DD protein levels were further increased compared to wildtype, Stmn2AA, and Stmn2CS (Figure C&D). Fluorescence microscopy confirmed that all Stmn2-Venus constructs were present in distal axons at relative levels consistent with western blot analysis (Figure 1E). As expected, palmitoylation-dead versions of Stmn2-Venus displayed a diffuse localization pattern compared to wildtype Stmn2 and Stmn2AA. Overall, these data demonstrate that palmitoylation is required for Stmn2-mediated protection of severed axons.

### The tubulin binding region contributes to enrichment of Stmn2 in axons

Stmn2 possesses a membrane targeting domain (MTD), a Stathmin-like and proline rich domain (designated together here as PrD), and tandem tubulin binding regions (TBR) (Figure 2A) (26). We demonstrated a functional role for Stmn2 palmitoylation in the MTD for suppressing axon degeneration (Figure 1) however a role for the Stmn2 TBR has not been evaluated. To address this question, we expressed a version of Stmn2AA lacking the TBR (ΔTBR-AA) and measured the kinetics of axon degeneration after axotomy. Expression of ΔTBR-AA accelerated axon degeneration compared to a vector control (Figure 2A). Notably, preventing palmitoylation of this Stmn2 construct (ΔTBR-DD) resulted in axon degeneration kinetics that matched the control.

**Figure 2.**
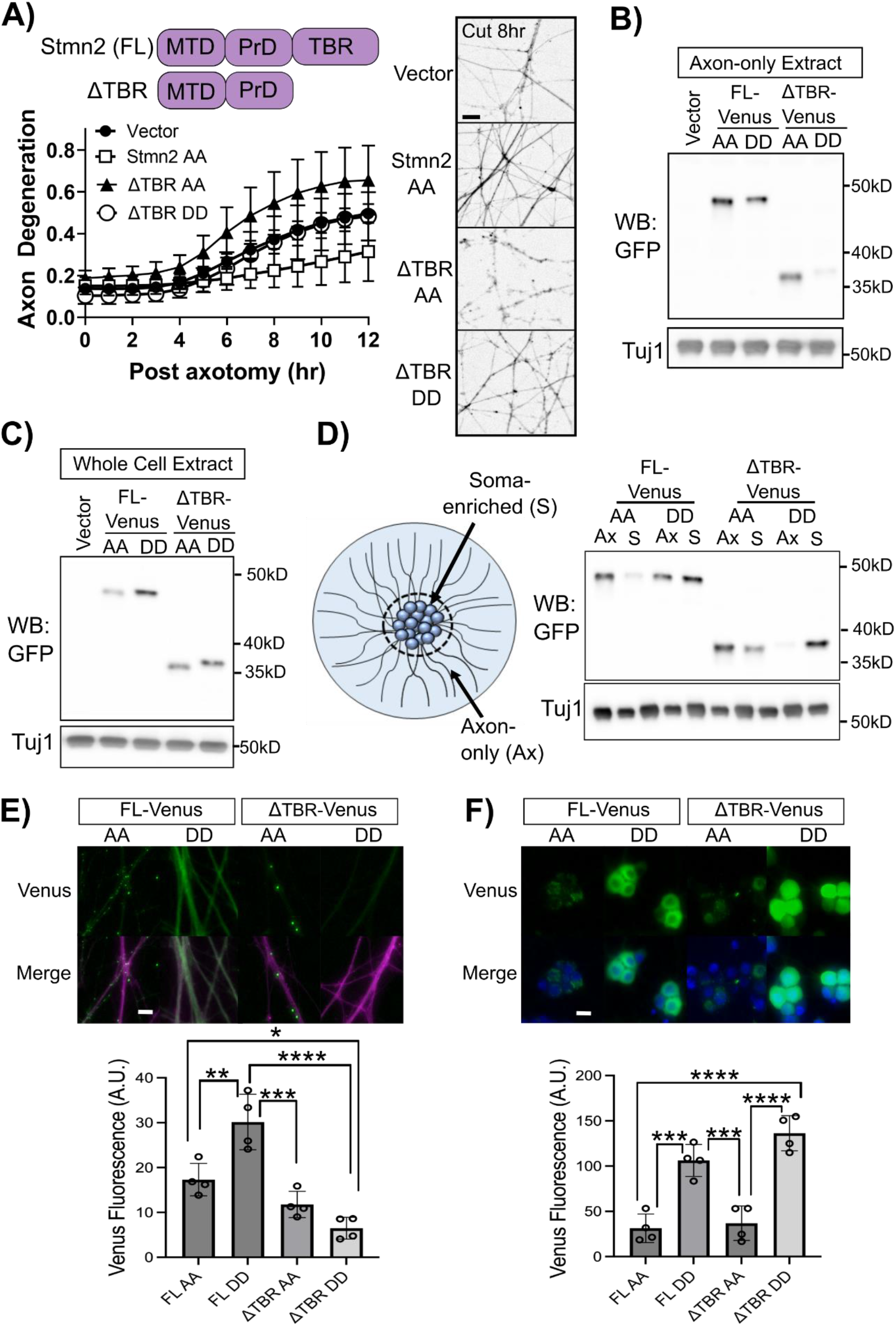
The tubulin binding region participates in Stmn2 localization and axon protection. *(A)* Domain structure of Stmn2 identifying the membrane targeting domain (MTD), Stathmin-like/Proline-rich domain (PrD) and tubulin binding region (TBR). We monitored the kinetics of axon degeneration after axotomy in the presence of the indicated Stmn2 constructs (full length or ΔTBR; AA S62,73A; DD C22,24S and S62,73A) (n = 4). Representative images of distal axons labeled with myristoylated mRuby3 at 8hr post axotomy shown to the right. Scale bar = 5μm. Western immunoblot detection of Venus-tagged Stmn2 constructs from *(B)* axon-only extracts or *(C)* whole cell extracts. *(D)* Axon-only and soma-enriched fractions were collected from the same culture expressing the indicated Stmn2-Venus construct and detected together by western immunoblot. *(E)* Fluorescence microscopy of distal axons from DRG sensory neurons expressing the indicated Stmn2-Venus construct and myristoylated mRuby3 to label axons (magenta). Quantification of Venus fluorescence shown below (n = 4). *(F)* Fluorescence microscopy of Venus expression in the soma of DRG sensory neurons and merged image with nuclei labeled with Hoechst. Quantification of fluorescence shown below (n = 4). For (*E*) and (*F*), means were compared with one-way ANOVA and statistical results from Bonferroni post-hoc tests shown in the graphs. Error bars represent +/-1 STD (* p< 0.05, ** p< 0.01, *** p< 0.001, **** p< 0.0001). Scale bar = 20μm.

Since expressing ΔTBR-AA did not suppress axon degeneration we wanted to confirm Stmn2 fragments used in Figure 2A were localizing to axon segments. We observed equivalent levels of ΔTBR-AA compared to full length forms of Stmn2 in axon-only extracts. However, axonal levels of ΔTBR-DD were reduced compared to full length Stmn2AA, Stmn2DD, and ΔTBR-AA (Figure 2B). When we examined whole cell extracts, ΔTBR-DD protein levels were comparable to other Stmn2 constructs (Figure 2C). Based on these findings, we speculated that ΔTBR-DD was retained in the soma. Axon-only and soma-enriched protein fractions were isolated from the same culture and directly compared (Figure 2D). Stmn2-AA and ΔTBR-AA were predominantly detected in the axon-only fraction while Stmn2-DD was detected in both axon-only and soma-enriched fractions, consistent with a predicted role for palmitoylation in vesicle-association and anterograde transport. However, ΔTBR-DD was largely restricted to the soma-enriched fraction (Figure 2D). Fluorescence microscopy of the Venus-tagged Stmn2 fragments were consistent with our biochemical studies (Figure 2E&F). Levels of Venus-tagged ΔTBR-DD were reduced in the axon compared to other Venus-tagged Stmn2 constructs yet this protein was readily detected in the soma. Collectively, these findings indicate that both palmitoylation and tubulin binding contribute to Stmn2 axonal localization. Moreover, the TBR is required for Stmn2-mediated axon protection after axotomy.

### The membrane targeting domain and proline rich domain of Stmn2 are required for axon protection

In contrast to the other Stathmins, Stmn1 does not possess a membrane targeting domain and is not palmitoylated (27) (Figure 3A). Overexpression of wildtype, human, Stmn1 did not suppress axon degeneration 10 hours post-axotomy (Figure 3B). We generated alanine substitutions in Stmn1 at serine 25 and serine 38 (Stmn1AA), which are phosphorylated by JNK (28) and correspond to the alanine substitutions in Stmn2AA (29). In contrast to Stmn2AA, overexpression of Stmn1AA did not suppress axotomy-induced axon degeneration (Figure 3B).

**Figure 3.**
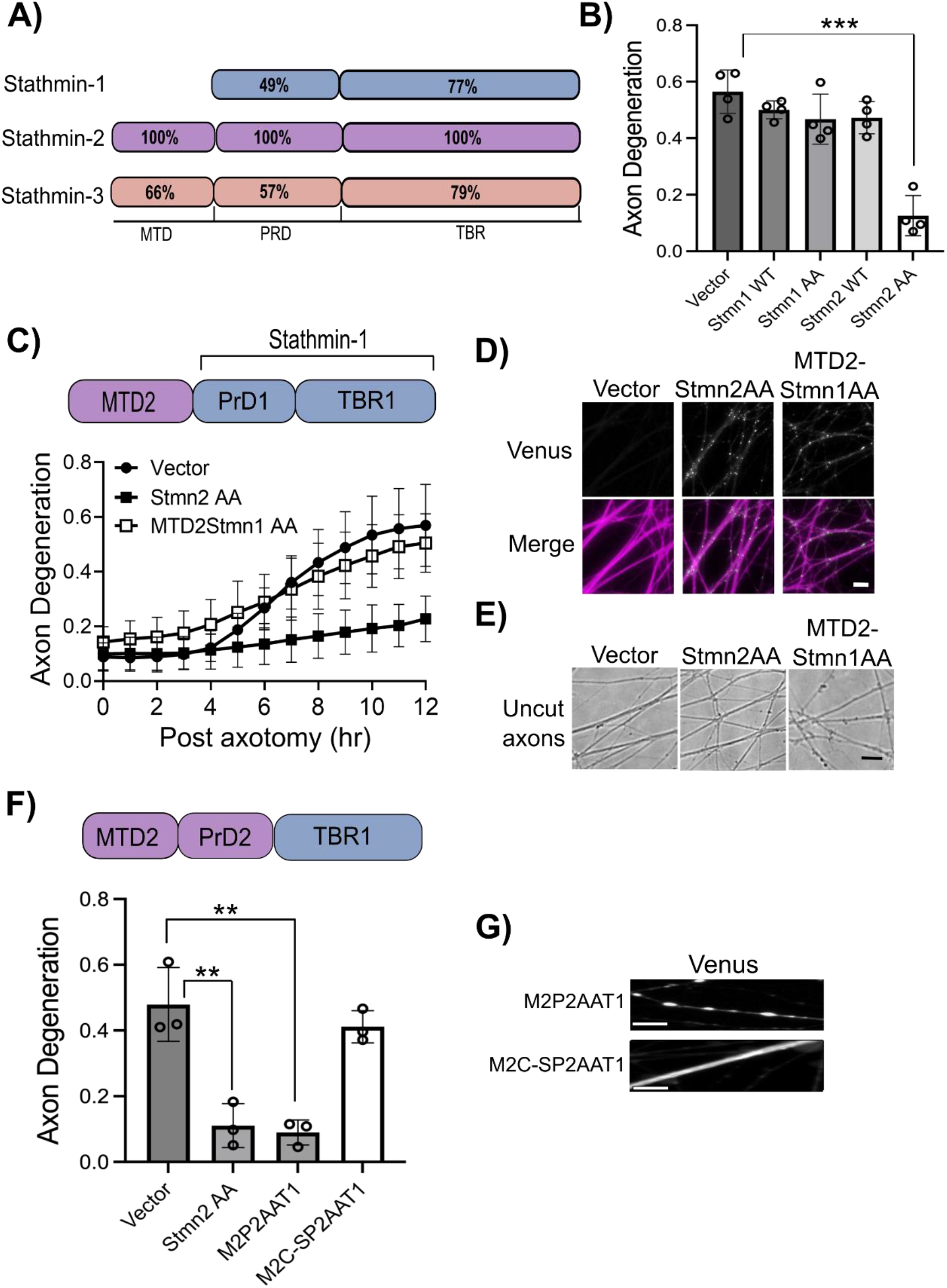
The Stmn1 tubulin binding region functionally compensates for the tubulin binding region of Stmn2 in axon protection. *(A)* Domain structures of Stmn1, Stmn2, and Stmn3 with percent homology displayed relative to Stmn2. *(B)* Degeneration of transected axons from DRG sensory neurons expressing the indicated Stathmin construct (n = 4, unpaired t-test). *(C)* Kinetics of axon degeneration 12hrs after axotomy from DRG sensory neurons expressing Stmn2AA or MTD2-Stmn1AA compared to an empty vector (n = 4). *(D)* MTD2-Stmn1AA-Venus is localized to puncta and observed in axons. *(E)* Phase contrast images of uncut, distal axons. Axonal blebbing is apparent in DRGs expressing MTD2-Stmn1AA-Venus. Scale bar in *(D)* and *(E)* equals 15 μm. *(F)* Axon degeneration in DRG sensory neurons 10hr post axotomy expressing Venus-tagged Stmn2AA or a Stmn2 AA chimera with the Stmn1 tubulin binding region (M2P2AAT1). We also evaluated axon degeneration in the presence of a M2P2AAT1 construct with mutations that impair palmitoylation (M2C-P2AAT1). Each construct was compared to the empty vector control (n = 4, unpaired t-test). *(G)* Venus-tagged Stmn2 chimeras were localized to distal axons. Scale bar = 5μm. Error bars represent +/-1 STD (* < 0.05, ** < 0.01, *** < 0.001, **** < 0.0001).

Since palmitoylation is required for Stmn2-mediated axon protection, we predicted that adding the Stmn2 membrane targeting domain (MTD) to Stmn1AA would endow this Stmn1 construct with axon protective activity. We generated a construct wherein the MTD of Stmn2 (aa1-40) was placed on the N-terminus of full-length Stmn1AA (MTD2Stmn1AA). However, overexpression of MTD2Stmn1AA did not affect the kinetics of axon degeneration over 12 hours post-axotomy (Figure 3C). MTD2Stmn1AA was localized to axon segments at similar levels to Stmn2AA (Figure 3D); however, overexpression of MTD2Stmn1AA resulted in baseline toxicity prior to axotomy, as indicated by axonal blebbing (Figure 3E), and confounding our ability to evaluate this question.

We next generated a construct that incorporates both the MTD and proline rich domain (PrD) of Stmn2 (containing alanine substitutions at Ser62 and Ser73) with the tubulin-binding domain of Stmn1 (M2P2AAT1). Overexpression of M2P2AAT1 suppressed axon degeneration to a similar extent as Stmn2AA (Figure 3F). Based on our findings in Figure 1, we predicted that activity of this chimera would be dependent on palmitoylation. Indeed, cysteine to serine substitutions that block palmitoylation (M2C-SP2AAT1) abrogated axon-protective activity (Figure 3F). Fluorescence microscopy confirmed that both constructs are present in distal axon segments and preventing palmitoylation results in diffuse localization (Figure 3G). Consequently, the Stmn2 MTD and PrD can function with the Stmn1 TBR to suppress axon degeneration.

### Stmn3 possess axon protective activity that is palmitoylation-dependent

Within the Stathmin family Stmn2 shares the most homology with Stmn3. We predicted that interfering with JNK-phosphorylated serines in the Stmn3 PrD would generate an axon-protective protein. The PrD of Stmn2 and Stmn3 share 57% homology (Figure 4A) however the Stmn3 PrD is phosphorylated at additional serines in this region by different kinases (30). We generated serine to alanine replacements in numerous residues previously identified as substrates for phosphorylation. These Stmn3 variants were expressed in DRGs and the degeneration of severed axons measured 10hr post axotomy. *In vitro* studies with Stmn3 identify Ser60 as the predominant JNK target in and Ser73 corresponds to the serine in Stmn2 predominantly phosphorylated by JNK (29). An alanine replacement in Ser60,73 (Stmn3 AA1) conferred very modest axon protection in when overexpressed (Figure 4B&C). We generated an additional alanine replacement at Ser62 (Stmn3 AAA) which corresponds to a modified serine in Stmn2AA based on sequence alignment, however this Stmn3 variant performed comparably to Stmn3 AA1 in axon protection.

**Figure 4.**
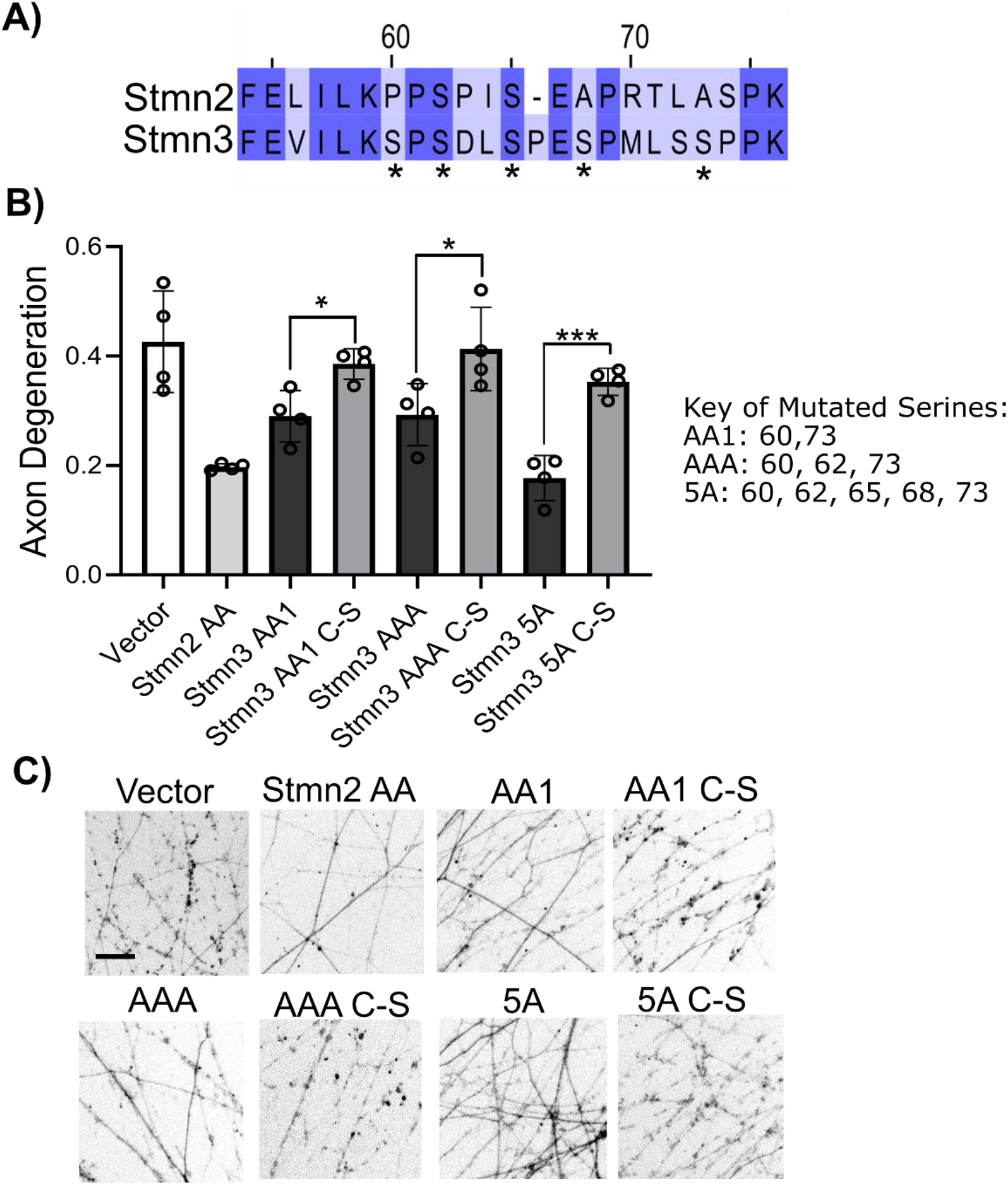
Stmn3 displays axon protective effects that are palmitoylation-dependent. *(A)* Sequence alignment of the proline rich (PrD) from Stmn2 and Stmn3. Asterisks identify serine residues in Stmn3 that are modified in this study. *(B)* Degeneration of transected axons from DRG sensory neurons expressing the indicated Stathmin constructs. Specific Stmn3 alanine-replacement constructs are described to the right. Each Stmn3 construct was also generated with amino acid substitutions that impair palmitoylation indicated by a C-S. Phosphorylation-dead constructs are compared with its matched palmitoylation-dead construct (n = 4, unpaired t-test). *(C)* Representative images of distal axons 10 hours post axotomy. Error bars represent +/-1 STD (* < 0.05, ** < 0.01, *** < 0.001, **** < 0.0001). Scale bar = 5μm.

Since there are additional putative phosphorylation sites in the Stmn3 PrD, we generated a Stmn3 variant with alanine replacements at five serines in the PrD (Stmn3 5A). Expressing Stmn3 5A resulted in axon protection that was comparable to Stmn2 AA in severed axons (Figure 4B&C). Stmn3 is palmitoylated (17,31) and we predicted that interfering with this post-translational modification would reverse the axon protective activity of these Stmn3 proteins. We generated cysteine to serine substitutions in the Stmn3 MTD of all three Stmn3 variants. Cysteine to serine substitutions in the Stmn3 MTD that prevent palmitoylation also suppressed the axon-protective activity of these Stmn3 alanine variants (Figure 4B&C). These data reinforce the functional contribution of palmitoylation to Stathmin-mediated axon protection.

### Stmn2 and Stmn3 comigrate in sensory neuron axons

The MTDs from Stmn2 and Stmn3 share considerable homology (66% identity) (Figure 5A) and undergo palmitoylation at two cysteine residues within this domain. To evaluate whether these domains can functionally compensate for one another, we swapped the MTD of Stmn2 with the MTD of Stmn3 in the presence of the axon protective Stmn2 AA substitution (M3P2AAT2). Expression of this chimeric protein suppressed axon degeneration to a similar extent as Stmn2-AA at 10 hours post axotomy (Figure 5B). Blocking palmitoylation with cysteine to serine substitutions in the Stmn3 MTD (M3C-P2AAT2) reversed this axon protective activity. Wildtype and palmitoylation-dead versions of this chimera were both localized in axons; however, the palmitoylation-dead version was diffuse (Figure 5C).

**Figure 5.**
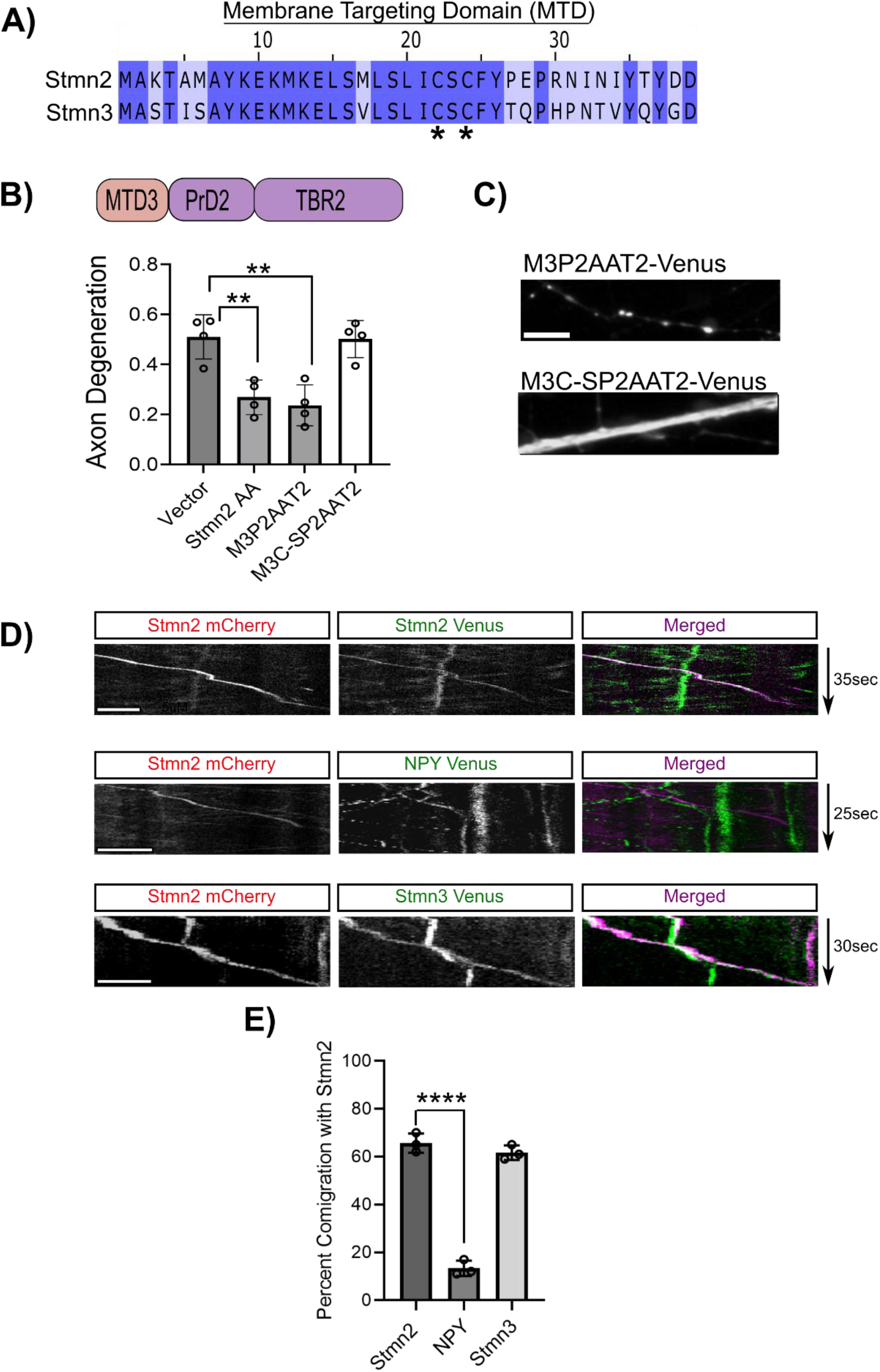
Stmn2 and Stmn3 comigrate in sensory neuron axons. *(A)* Sequence alignment of the membrane targeting domain (MTD) of Stmn2 and Stmn3. Asterisks indicate palmitoylation sites. *(B)* Degeneration of transected axons from DRG sensory neurons 10 hours post-axotomy expressing Venus-tagged Stmn2 AA or indicated Stmn2AA chimera with the MTD of Stmn3 (M3P2AAT2). We also evaluated axon degeneration in the presence of chimeras with mutations that impair palmitoylation (M3C-SP2AAT2). Each construct is compared to the empty vector control (n = 4, unpaired t-test). *(C)* Localization of Venus-tagged M3P2AAT2 and M3C-SP2AAT2 in axon segments. Scale bar represents 5μm *(D)* Representative kymographs from live-imaging studies following comigration between Stmn2-mCherry puncta with either Stmn2-Venus (top panels), NPY-Venus (middle panels), or Stmn3-Venus (bottom panels) in axon segments. Grayscale kymographs are included for individual channels with merged kymograph to show puncta comigration. Scale bars represent 5μm. *(E)* Quantification of comigration with Stmn2 (n = 3, unpaired t-test). Error bars represent +/-1 STD (* < 0.05, 723 ** < 0.01, *** < 0.001, **** < 0.0001).

Since the MTD of Stmn3 functionally compensates for the MTD of Stmn2, we hypothesized that Stmn2 and Stmn3 are targeted to the same vesicle population. We transduced differentially tagged versions of Stmn2 and Stmn3 constructs into DRG sensory neurons and used live imaging to track comigration. To assess maximal comigration we expect from our assay, we first co-transduced mCherry and Venus-tagged versions of Stmn2 and measured the rate of comigration in axon segments. From this analysis, we observed 65% comigration between Stmn2-containing particles (Figure 5D &E). Next, we evaluated comigration between Stmn2-mCherry and Stmn3-Venus particles in axon segments. We observed 62% comigration between Stmn2-mCherry and Stmn3-Venus labeled particles, which is comparable to our comigration measurements with differentially tagged Stmn2 proteins. To assess the specificity of this assay, we measured comigration between Stmn2-mCherry with Neuropeptide Y (NPY). NPY is trafficked through secretory vesicles that are predicted to have lower rates of comigration with Stmn2-mCherry compared to Stmn3. We observed 13% comigration between Stmn2-mCherry and NPY-EGFP, suggesting minimal comigration between these two proteins (Figure 5D&E). Therefore, Stmn2 and Stmn3 comigrate on the same vesicle population and the membrane targeting domains are functionally equivalent in axon maintenance.

### The membrane targeting domain of Stmn2 promotes DLK-dependent degradation

Our experiments identify a critical role for membrane localization and palmitoylation in Stmn2 axon protective function in severed axons. Palmitoylation also contributes to regulated turnover of Stmn2 protein (25). Stmn2 is a short-lived protein and axon transection leads to depletion of Stmn2 protein from severed axons in DRG sensory neurons (32). To determine if this property is unique to Stmn2, we evaluated whether Stmn1 and Stmn3 are depleted in axons after axotomy. At four hours post axotomy, protein levels of Stmn2 and Stmn3 were reduced below 10% of uncut control (Figure 6A). Stmn1 protein levels were unchanged compared to uncut control.

**Figure 6.**
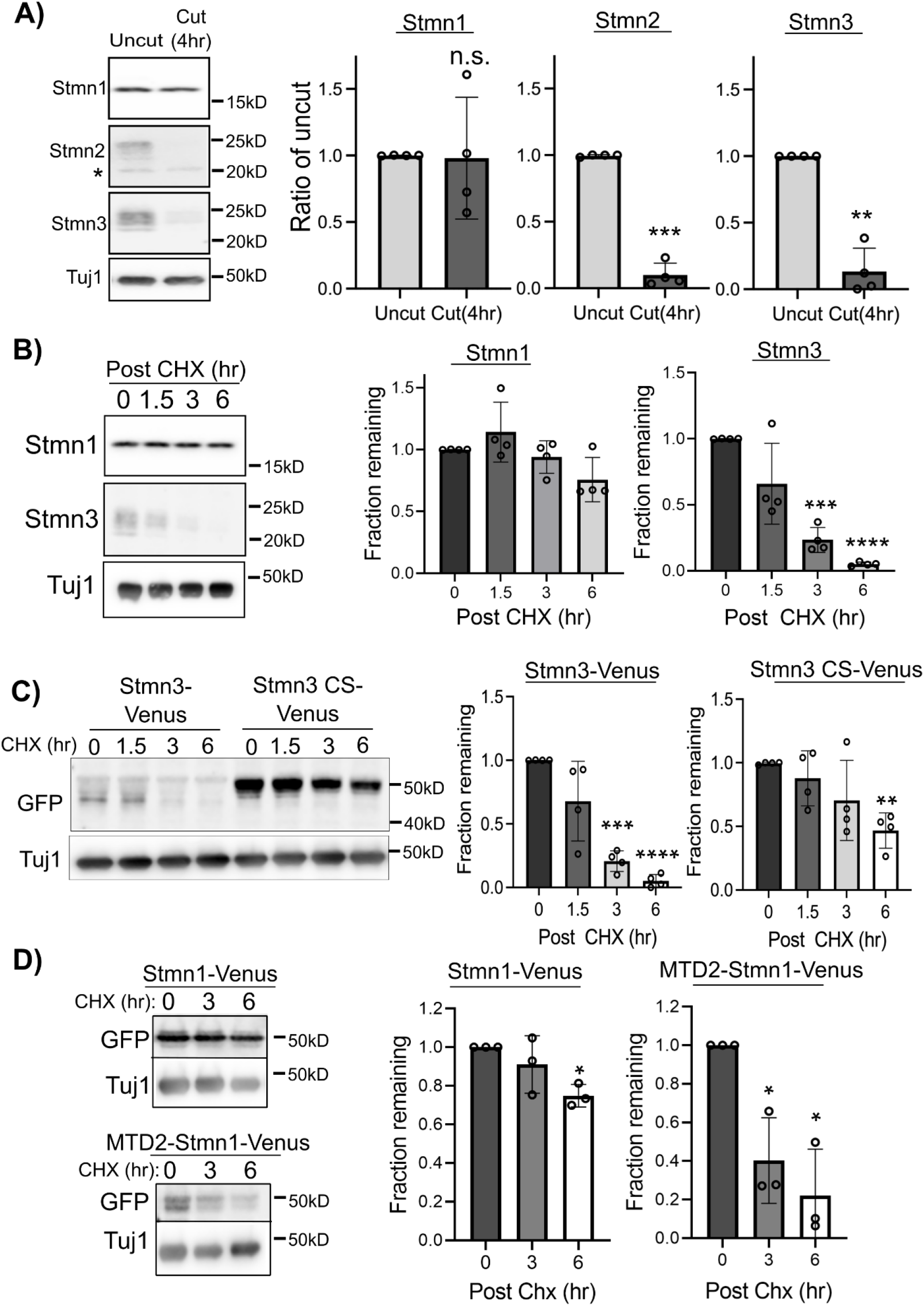
Membrane associated Stathmins are targeted for degradation. *(A)* Western immunoblot analysis of endogenous Stmn1, Stmn2, and Stmn3 from intact or severed axons 4hr post cut (n = 4, unpaired Welch’s t-test; n.s. = not significant). *(B)* Protein turnover of endogenous Stmn1 and Stmn3 following addition of cycloheximide (CHX) to inhibit protein synthesis for the indicated times. Each time point was compared to the 0hr control (n=4, unpaired Welch’s t-test). *(C)* Protein turnover of Venus-tagged Stmn3 constructs (CS; C22,24S) detected via western immunoblotting following CHX addition. Each time point was compared to the 0hr control (n = 4, unpaired Welch’s t-test). *(D)* Protein turnover of Venus-tagged Stmn1 (Stmn1-Venus) and membrane targeted Stmn1 (MTD2-Stmn1-Venus) detected via western immunoblotting following addition of CHX. Each time point was compared to the 0hr control (n=4, unpaired Welch’s t-test). Representative western blots shown to the left. Error bars represent +/-1 STD (* < 0.05, 737 ** < 0.01, *** < 0.001, **** < 0.0001).

We next evaluated the half-life of endogenous Stmn1 and Stmn3 in axon segments. We inhibited protein synthesis with cycloheximide (CHX) and monitored loss of endogenous Stmn1 and Stmn3 in axon-only extracts by western immunoblotting. The half-life of Stmn3 in axons was approximately two hours (Figure 6B), similar to the half-life of Stmn2. In contrast, Stmn1 protein levels were unchanged at three hours post CHX treatment and reduced by only 20% after six hours of CHX treatment. We examined whether palmitoylation contributes to Stmn3 turnover as observed in Stmn2 (25). We compared protein turnover rates between wildtype Stmn3-Venus with a version that is not palmitoylated (Stmn3CS-Venus). The half-life of Stmn3CS-Venus was extended to six hours (Figure 6C) supporting a role for palmitoylation in protein turnover of both Stmn2 and Stmn3 in axons.

Stmn1 is not associated with membranes and displays greater stability in axon segments compared to Stmn2 and Stmn3. We attached the Stmn2 MTD to the N-terminus of Stmn1 to ascertain whether the Stmn2 MTD could confer instability on another Stathmin. Stmn1-Venus or MTD2-Stmn1-Venus were expressed in DRGs and protein turnover evaluated in axon-only extracts after CHX treatment. Consistent with our measurements of endogenous Stmn1, Venus-tagged Stmn1 decreased approximately 20% after six hours CHX treatment (Figure 6D). On the other hand, MTD2-Stmn1-Venus protein levels decreased 80% by six hours post-CHX addition indicating the Stmn2 MTD can confer instability on another Stathmin.

MAPK signaling via the DLK-JNK pathway promotes regulated turnover of palmitoylated Stmn2 (25). Whether DLK signaling regulates axonal levels of other Stathmin proteins is not known. Acutely blocking DLK or JNK activity with small molecule inhibitors (1μM GNE-3511 and 5μM JNK inhibitor VIII) increased steady state, axonal levels of endogenous Stmn3 yet did not affect endogenous Stmn1 (Figure 7A & B). Since adding the Stmn2 MTD to Stmn1 accelerated its turnover, we evaluated whether the Stmn2 MTD confers sensitivity to DLK signaling. For these experiments, expression constructs were generated with a Flag epitope. GNE-3511 treatment did not affect Stmn1-Flag levels in axon-only extracts. However, GNE-3511 treatment did induce a significant increase in Stmn2-Flag, and MTD2-Stmn1-Flag protein levels (Figure 7C&D). As a complementary approach, we used Venus-tagged versions of these constructs to quantify changes in Venus’s fluorescence intensity in distal axon segments (Figure 7E). Stmn1-Venus fluorescence intensity levels were not affected by GNE-3511. However, GNE-3511 treatment induced a significant increase in Stmn2-Venus and MTD2-Stmn1-Venus fluorescence intensity (Figure 7F).

**Figure 7.**
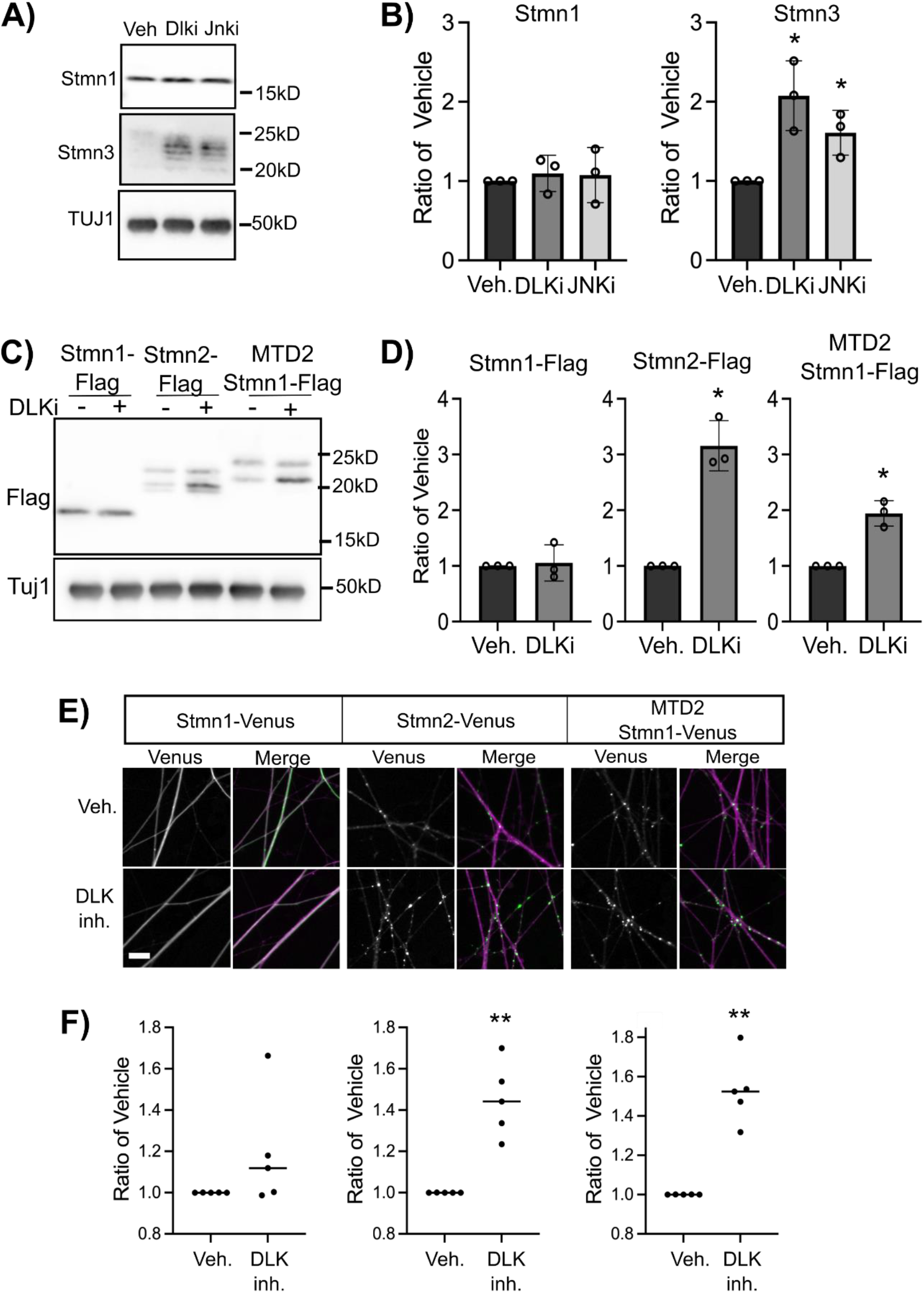
Membrane association targets Stathmins for DLK-dependent degradation. *(A)* Representative western blots of endogenous Stmn1 and Stmn3 following acute treatment with DLK and JNK inhibitors for 4hr with quantification shown in *(B)*, drug treatment was compared to vehicle control (n = 4, unpaired Welch’s t-test) *(C)* Representative western blot of Flag-tagged Stmn1, Stmn2 and membrane targeted Strmn1 (MTD2 Stmn1) following treatment with vehicle or DLKi (4hr), quantification is shown in *(D)* (n = 4, unpaired Welch’s t-test). *(E)* Representative images of Venus-tagged Stmn1, Stmn2, and MTD2 Stmn1 treated with vehicle or DLKi (4hr). Merge images show overlay between Venus and myristoylated mRuby3 (magenta) used to label axons. Quantification is shown in *(F)* (n = 4, unpaired Welch’s t-test). Error bars represent +/-1 750 STD (* < 0.05, ** < 0.01, *** < 0.001, **** < 0.0001). Scale bar = 20μm.

### The Stmn2 MTD is sufficient to promote vesicle comigration and DLK-dependent degradation

Attaching the MTD from Stmn2 to GFP relocalizes this fluorescent protein to Golgi as well as vesicles in the growth cones of hippocampal neurons (33). The authors of this study also observed co-localization between this GFP fusion and endogenous Stmn2. We attached the Stmn2 MTD to Venus and evaluated localization, comigration with Stmn2, and protein turnover in DRG sensory neurons. MTD2-Venus localized to puncta in the soma and axon, similar to full-length Stmn2 (Figure 8A&B). We also observed a stronger diffuse MTD2-Venus population in both axons and soma compared to Stmn2-Venus. Focusing on the population of axonal vesicles, we measured comigration between MTD2-Venus with full-length Stmn2-mCherry. We observed 58% comigration between MTD2-Venus with Stmn2-mCherry (Figure 8C), which is comparable to the level of comigration between differentially tagged forms of Stmn2 (Figure 5).

**Figure 8.**
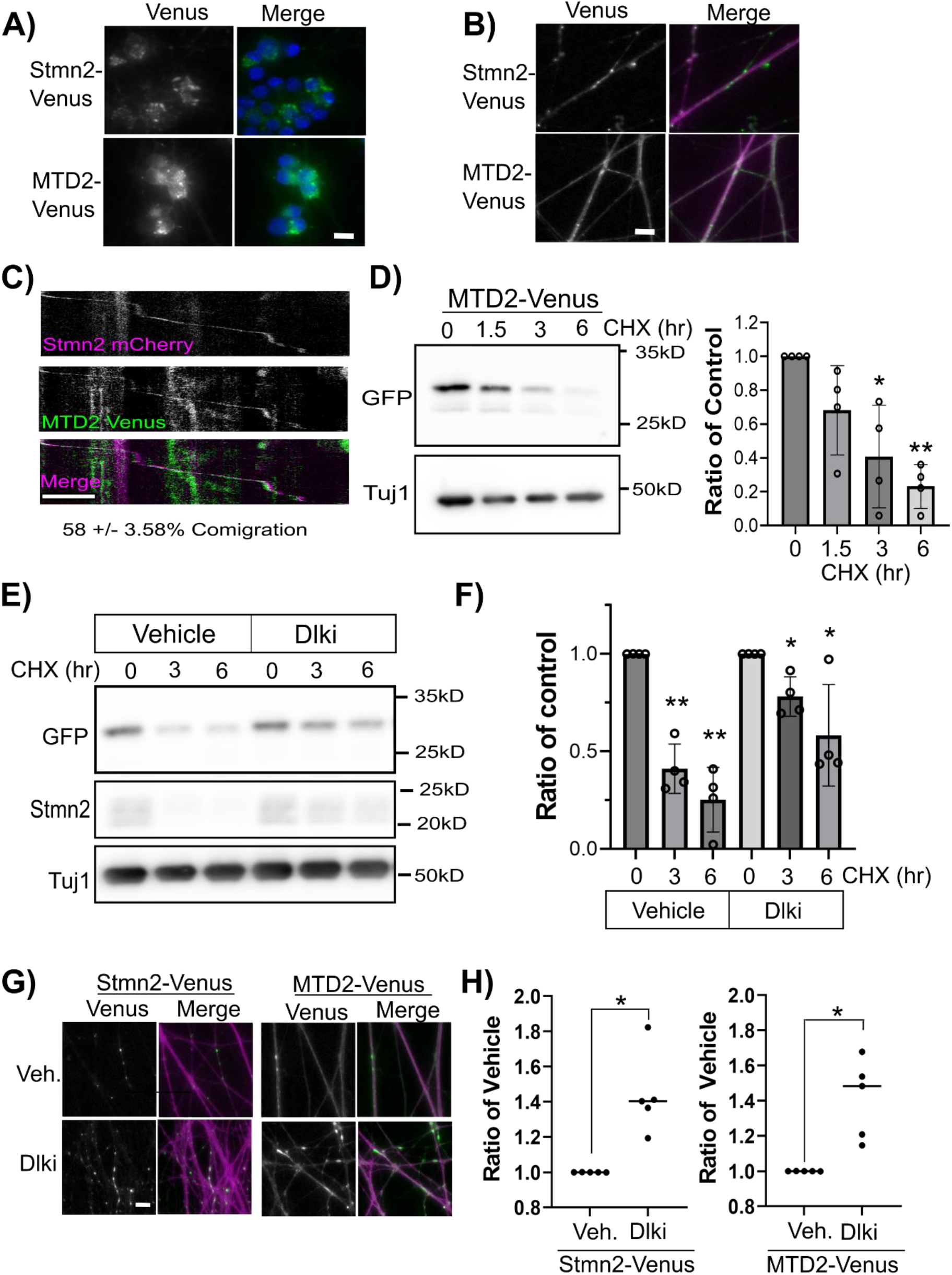
The membrane targeting domain is sufficient for vesicular targeting and regulation by DLK. Representative images of Venus-tagged full-length Stmn2 (Stmn2-Venus) and just the membrane targeting domain (MTD2-Venus). Images are merged with either Hoechst to stain for the cell body *(A)* or myristoylated mRuby3 to label axons *(B). (C)* Representative kymographs of Stmn2 mCherry and MTD2 Venus showing comigration (Scale bar = 5μm). Comigration of MTD2 with Stmn2 determined to be 58 +/-3.58% (n = 3, unpaired t-test). *(D)* Levels of the Venus-tagged membrane targeting domain of Stmn2 (MTD2) were detected via western immunoblotting following treatment with cycloheximide for the indicated time period. Each time point was compared to the 0hr control (n = 4, unpaired Welch’s t-test). Representative western blot shown to the left. *(E)* Representative western blot of Venus-tagged MTD2 following acute treatment with vehicle or DLKi, and treatment with cycloheximide at indicated time periods. Endogenous Stmn2 shown as a control for DLK treatment. Quantification shown in (*F*), each time point was compared to 0hr control (n = 4, unpaired Welch’s t-test; vehicle). *(G)* Representative images of Venus-tagged Stmn2 and MTD2 following treatment with vehicle or DLKi. With quantification shown in (*H*), DLKi compared to the vehicle (n = 4, unpaired Welch’s t-test). Error bars represent 770 +/-1 STD (* < 0.05, ** < 0.01, *** < 0.001, **** < 0.0001). Scale bar = 20μm.

Considering differential stability between Stmn2 and Stmn1 and the impact of adding the Stmn2 MTD to Stmn1, we next determined whether the Stmn2 MTD is sufficient to promote regulated degradation of a protein other than a Stathmin. We measured turnover of MTD2-Venus in axon-only extracts after CHX treatment. MTD2-Venus levels decreased by 30% after 1.5 hours CHX treatment and 80% by 6 hours (Figure 8D). Since Stmn2 degradation is regulated by DLK signaling, we next examined whether MTD2-Venus degradation was dependent on this pathway. In the presence of the DLK inhibitor GNE-3511, the turnover kinetics of the MTD2 were slowed, with only a 20% decrease after 3hr CHX treatment and 40% drop by 6hrs (Figure 8E&F). We also examined steady state levels in axons with fluorescence microscopy in response to DLK inhibition. Axonal MTD2-Venus levels increased to a comparable level as full length Stmn2-Venus after 4 hours GNE-3511 treatment. (Figure 8G&H). We conclude that the Stmn2 MTD promotes localization to a specific Stmn2-containing vesicle population and confers regulated degradation via the DLK signaling.

## DISCUSSION

Stathmins are a family of phosphoproteins with a long-appreciated function in microtubule dynamics. While heavily studied in the context of neurite outgrowth, Stathmins also contribute to axon maintenance. Stmn1^-/-^ mice develop late onset axonopathy (34) and loss of Stmn2 is linked to motor neuropathy in mice and humans (4-8). In particular, re-expressing Stmn2 in ALS-derived human motor neurons rescues axon outgrowth defects *in vitro* (4,5). Therefore, restoring Stmn2 protein in axon segments is a potential therapeutic opportunity however mechanisms required for Stmn2 axon protective activity are not clear. We demonstrate that palmitoylation is required for Stmn2-mediated axon protection. Notably, blocking Stmn2 palmitoylation did not prevent Stmn2 enrichment within the axon segment. Instead, tubulin interaction through the tubulin binding region also promotes Stmn2 localization to axons suggesting the relationship between Stathmins and the microtubule cytoskeleton is bidirectional. Therefore, Stathmins regulate microtubule dynamics by sequestering tubulin heterodimers while also using the microtubule network for transport into distal axon segments.

In contrast to other members of the Stathmin family, Stmn1 does not possess a membrane targeting domain and did not display inherent axon protective activity in severed axons. Relocating Stmn1 to a membrane using the Stmn2 MTD triggered spontaneous axonal blebbing and did not confer axon-protective activity. In contrast, introducing the Stmn2 MTD and PrD on the Stmn1 TBR was well-tolerated and generated an axon protective Stathmin protein. There might be intramolecular interactions between the MTD and PrD that influence Stathmin function in a cell. The PrD is an unstructured region present in all Stathmins yet also displays the highest divergence in amino acid sequence (11). Moreover, relocalizing Stmn1 to a vesicle might interfere with normal protein: protein interactions on the Stmn2 vesicle that lead to toxicity.

The identity of the Stmn2 vesicle population is not known. Stmn3 comigrates with Stmn2 in axon segments suggesting there are vesicles enriched with Stathmin proteins. Stmn3 is also phosphorylated by JNK (29) and alanine-substitutions in the Stmn3 PrD conferred partial axon-protective activity however the effects were noticeably modest compared to Stmn2AA. This difference might reflect unique functional properties in Stmn2 that contribute to axon maintenance and why loss of Stmn2 is sufficient to provoke axonopathy (7-9). The Stmn3 MTD can functionally compensate for the Stmn2 MTD and palmitoylation is required for Stmn3 axon-protective activity. Whether membrane association affects interaction between Stmn2 and tubulin heterodimers is not known. Palmitoylation is dispensable for tubulin binding *in vitro* however other residents on the Stmn2 vesicle could influence this interaction inside the cell. Stmn2 palmitoylation also contributes to its function in protein trafficking of Amyloid Precursor Protein and chromaffin (20,21,35). Moreover, Stmn2 regulates mitochondrial transport in axons (15) indicating a broader, still unclear function in organelle transport that could have significant relevance to axon maintenance and dysfunction in disease.

Membrane association is consistently connected to protein turnover of Stathmin proteins. Both Stmn2 and Stmn3 are short-lived proteins and degradation is regulated by DLK-JNK signaling. Alanine replacements in JNK-phosphorylated serines extends Stmn2 stability suggesting direct phosphorylation in the PrD promotes degradation (32). However, attaching the Stmn2 MTD alone to a reporter protein shortens half-life and confers sensitivity to DLK-JNK signaling even in the absence of the Stmn2 PrD. JNK-mediated phosphorylation of the Stmn2 MTD is not reported, yet cannot be ruled out. This MTD promotes comigration with Stmn2 on the same vesicle population and might be targeted for degradation by association. Interestingly, many proteins in the DLK-JNK signaling complex are palmitoylated, including DLK itself (36,37). DLK signaling is activated in response to microtubule dysfunction (38-40) raising the possibility that DLK is a conduit between the microtubule cytoskeleton and Stmn2 activity. Understanding how microtubules, DLK, and Stathmins communicate in this signaling circuit will generate important insight on mechanisms of axon maintenance.

## ACKNOWLEDGEMENTS

We thank members of the Summers lab for their helpful comments during preparation of this manuscript. This work was supported by a grant from the National Institutes of Health (RO1NS126191 to DWS). The content is solely the responsibility of the authors and does not necessarily reflect the official views of the National Institutes of Health.

